# SNPs associated with *HHIP* expression have differential effects on lung function in males and females

**DOI:** 10.1101/594457

**Authors:** KA Fawcett, M Obeidat, CA Melbourne, N Shrine, AL Guyatt, C John, J Luan, A Richmond, MR Moksnes, R Granell, S Weiss, M Imboden, S May-Wilson, P Hysi, TS Boutin, L Portas, C Flexeder, SE Harris, CA Wang, L Lyytikäinen, T Palviainen, RE Foong, D Keidel, C Minelli, C Langenberg, Y Bossé, Berge M van den, D Sin, K Hao, A Campbell, D Porteous, S Padmanabhan, BH Smith, D Evans, S Ring, A Langhammer, K Hveem, C Willer, R Ewert, B Stubbe, N Pirastu, L Klaric, PK Joshi, K Patasova, M Massimo, O Polasek, JM Starr, I Rudan, T Rantanen, K Pietiläinen, M Kähönen, OT Raitakari, GL Hall, PD Sly, CE Pennell, J Kaprio, T Lehtimäki, V Vitart, IJ Deary, D Jarvis, JF Wilson, T Spector, N Probst-Hensch, N Wareham, H Völzke, J Henderson, D Strachan, BM Brumpton, C Hayward, IP Hall, MD Tobin, LV Wain

**Affiliations:** Department of Health Sciences, University of Leicester, Leicester, LE1 7RH, UK; The University of British Columbia Centre for Heart Lung Innovation, St Paul’s Hospital, Vancouver, BC, Canada; MRC Epidemiology Unit, University of Cambridge School of Clinical Medicine, Cambridge, CB2 0QQ, UK; MRC Human Genetics Unit, Institute of Genetics and Molecular Medicine, University of Edinburgh, Western General Hospital, Crewe Road, Edinburgh, EH4 2XU, Scotland; K.G. Jebsen Center for Genetic Epidemiology, Department of Public Health and Nursing, NTNU, Norwegian University of Science and Technology, Norway; Integrative Epidemiology Unit, University of Bristol, Bristol, UK, BS8 2BN; Department of Functional Genomics, Interfaculty Institute for Genetics and Functional Genomics, University Medicine Greifswald, 17475 Greifswald, Germany; Swiss Tropical and Public Health Institute, Basel, Switzerland; University of Basel, Switzerland; Centre for Global Health Research, Usher Institute of Population Health Sciences and Informatics, University of Edinburgh, Teviot Place, Edinburgh, EH8 9AG, Scotland; The Department of Twin Research & Genetic Epidemiology, King’s College London, St Thomas’ Campus, Lambeth Palace Road, London, SE1 7EH; Population Health and Occupational Disease, National Heart and Lung Institute, Imperial College London, London, UK; Institute of Epidemiology, Helmholtz Zentrum München - German Research Center for Environmental Health, 85764 Neuherberg, Germany; Centre for Cognitive Ageing and Cognitive Epidemiology, University of Edinburgh, Edinburgh, UK, EH8 9JZ; Psychology, University of Edinburgh, Edinburgh, UK, EH8 9JZ; School of Medicine and Public Health, Faculty of Medicine and Health, The University of Newcastle; Department of Clinical Chemistry, Fimlab Laboratories, Tampere 33520, Finland; Department of Clinical Chemistry, Finnish Cardiovascular Research Center - Tampere, Faculty of Medicine and Health Technology, Tampere University, Tampere 33014, Finland; Department of Cardiology, Heart Center, Tampere University Hospital, Tampere 33521, Finland; Institute for Molecular Medicine FIMM, University of Helsinki, FI-00014 Helsinki, Finland; Telethon Kids Institute; School of Physiotherapy and Exercise Science, Faculty of Health Sciences, Curtin University; Institut Universitaire de Cardiologie et de Pneumologie de Québec, Québec, Quebec, Canada; University of Groningen, University Medical Center Groningen, Department of Pulmonology, GRIAC Research Institute, University of Groningen, Groningen, The Netherlands; Respiratory Division, Department of Medicine, University of British Columbia, Vancouver, BC, Canada; Department of Genetics and Genomic Sciences, Icahn School of Medicine at Mount Sinai, New York, NY, USA; Centre for Genomic and Experimental Medicine, Institute of Genetics and Molecular Medicine, Western General Hospital, EH4 2XU, UK; British Heart Foundation Glasgow Cardiovascular Research Centre, Institute of Cardiovascular and Medical Sciences, College of Medical, Veterinary and Life Sciences, University of Glasgow, Glasgow G12 8TA, UK; Division of Population Health Sciences, Ninewells Hospital and Medical School, University of Dundee, Dundee, DD1 9SY, UK; Population Health Sciences Bristol Medical School University of Bristol, Bristol, UK, BS8 2BN; Department of Public Health and Nursing, Faculty of Medicine and Health Sciences, NTNU, Norwegian University of Science and Technology, Trondheim, Norway; Department of Biostatistics and Center for Statistical Genetics, University of Michigan, Ann Arbor, USA; Department of Internal Medicine, University of Michigan, Ann Arbor, MI, US; Department of Human Genetics, University of Michigan, Ann Arbor, USA; Department of Internal Medicine B, Cardiology, Pneumology, Infectious Diseases, Intensive Care Medicine, University Medicine Greifswald, 17475 Greifswald, Germany; Faculty of Medicine, University of Split, Split, Croatia; Alzheimer Scotland Research Centre, University of Edinburgh, Edinburgh, UK, EH8 9JZ; Faculty of Sport and Health Sciences, Gerontology Research Center, University of Jyväskylä; Obesity Research Unit, Research Program for Clinical and Molecular Metabolism, Faculty of Medicine, University of Helsinki, FI-00014 Helsinki, Finland; Obesity Centre, Abdominal Centre, Helsinki University Hospital and University of Helsinki, FI-00029 Helsinki, Finland; Department of Clinical Physiology, Tampere University Hospital, Tampere 33521, Finland; Department of Clinical Physiology, Finnish Cardiovascular Research Center - Tampere, Faculty of Medicine and Health Technology, Tampere University, Tampere 33014, Finland; Centre for Population Health Research, University of Turku and Turku University Hospital, Finland; Research Centre of Applied and Preventive Cardiovascular Medicine, University of Turku, Finland; Department of Clinical Physiology and Nuclear Medicine, Turku University Hospital, Turku, Finland; Children’s Health and Environment Program, The University of Queensland; Department of Public Health, University of Helsinki, FI-00014, Helsinki, Finland; MRC-PHE Centre for the Environment and Health, London, UK; Intitute for Community Medicine, University Medicine Greifswald, 17487 Greifswald, Germany; Population Health Research Institute, St George’s, University of London, London SW17 0RE, UK; Clinic of Thoracic and Occupational Medicine, St. Olavs Hospital, Trondheim University Hospital; Division of Respiratory Medicine and NIHR-Nottingham Biomedical Research Centre, University of Nottingham; National Institute for Health Research, Leicester Respiratory Biomedical Research Centre, Glenfield Hospital, Leicester, LE3 9QP, UK

## Abstract

Adult lung function is highly heritable and 279 genetic loci were recently reported as associated with spirometry-based measures of lung function. Though lung development and function differ between males and females throughout life, there has been no genome-wide study to identify genetic variants with differential effects on lung function in males and females. Here, we present the first genome-wide genotype-by-sex interaction study on four lung function traits in 303,612 participants from the UK Biobank. We detected five SNPs showing genome-wide significant (P<5 × 10^−8^) interactions with sex on lung function, as well as 21 suggestively significant interactions (P<1 × 10^−6^). The strongest sex interaction signal came from rs7697189 at 4:145436894 on forced expiratory volume in 1 second (FEV_1_) (P = 3.15 × 10^−15^), and was replicated (P = 0.016) in 75,696 individuals in the SpiroMeta consortium. Sex-stratified analyses demonstrated that the minor (C) allele of rs7697189 increased lung function to a greater extent in males than females (untransformed FEV_1_ β = 0.028 [SE 0.0022] litres in males vs β = 0.009 [SE 0.0014] litres in females), and this effect was not accounted for by differential effects on height, smoking or age at puberty. This SNP resides upstream of the gene encoding hedgehog-interacting protein (*HHIP*) and has previously been reported for association with lung function and *HHIP* expression in lung tissue. In our analyses, while *HHIP* expression in lung tissue was significantly different between the sexes with females having higher expression (most significant probeset P=6.90 × 10^−6^) after adjusting for age and smoking, rs7697189 did not demonstrate sex differential effects on expression. Establishing the mechanism by which *HHIP* SNPs have different effects on lung function in males and females will be important for our understanding of lung health and diseases, such as chronic obstructive pulmonary disease (COPD), in both sexes.

## Introduction

Measures of lung function, including forced expiratory volume in 1 second (FEV_1_) and forced vital capacity (FVC), are used to determine diagnosis and severity of chronic obstructive pulmonary disease (COPD). COPD refers to a group of complex lung disorders characterised by irreversible (and usually progressive) airway obstruction, and is projected to be the third leading cause of death globally in 2020 (1). The major risk factor for COPD is smoking, but other environmental and genetic factors have been identified.

Physiological lung development and function differ throughout life between males and females. Female foetuses have smaller airways and fewer bronchi, but their lungs mature faster and they produce surfactant earlier than lungs of male foetuses (2). During childhood, females have smaller lungs compared to height-matched males, but have a higher flow rate per lung volume, perhaps reflecting airway growth lagging behind lung growth in males (3). In adulthood, females have smaller diameter airways, fewer alveoli, and smaller lung volumes and diffusion surfaces compared to males (4). However, there is some evidence to suggest that age-related decline in lung function is slower in females (5). As well as these anatomical differences, it is known that sex hormones can influence lung structure and function throughout life but the mechanisms are not well understood (5, 6).

The incidence and presentation of lung diseases such as COPD also exhibit sex disparity. Traditionally viewed as a disease of older males, COPD has been increasing in prevalence amongst females over the last two decades. It has been reported that females are more vulnerable to environmental risk factors for COPD and are over-represented amongst sufferers of early-onset severe COPD (7, 8). Females are also more likely to present with small airway disease whereas males are more likely to develop emphysematous phenotype. Moreover, females report more frequent and/or severe exacerbations of respiratory symptoms than males and higher levels of dyspnea and cough (7).

In a recent paper, 279 genetic loci were reported as associated with lung function traits, but these only explain a small proportion of the heritability (9). One possible source of hidden heritability is the interaction between genetic factors and biological sex on lung function traits. A genome-wide genotype-by-sex interaction study in three studies comprising 6260 COPD cases and 5269 smoking controls found a putative sex-specific risk factor for COPD in the *CELSR1* gene, a region not previously implicated in COPD or lung function (10). However, having sufficient statistical power to reproducibly detect genotype-by-sex interactions will require much larger sample sizes. Statistical power can also be enhanced by using quantitative lung function traits as outcomes but we are not aware of any genome-wide genotype-by-sex interaction studies on lung function traits. Understanding the role of sex in lung function and COPD will be important for developing therapeutics that work for both males and females (11).

In this study, we tested for an interaction effect of 7,745,864 variants and sex on FEV_1_, FEV_1_/FVC, FVC and PEF in 303,612 individuals from the UK Biobank resource. We sought replication of our findings in 75,696 independent individuals from the SpiroMeta consortium. To our knowledge this is the first genome-wide sex-by-genotype interaction study on lung function traits and the largest sex-by-genotype interaction study to focus on COPD-related outcomes.

## Materials and Methods

### UK Biobank participants

The UK Biobank is described here: http://www.ukbiobank.ac.uk. It comprises over 500,000 volunteer participants aged 40-69 years at time of recruitment, with demographic, lifestyle, clinical and genetic data (12). Individuals were selected for inclusion in this study if (i) they had no missing data for sex, age, height, and smoking status, (ii) their spirometry data passed quality control, as described previously (9), (iii) they had genome-wide imputed genetic data, (iv) they were of genetically determined European ancestry, and (v) they were not first-or second-degree relatives of any other individual included in the study. In total, 303,612 individuals met these criteria (**Supplementary Table 1**).

Participants’ DNA was genotyped using either the Affymetrix Axiom^®^ UK BiLEVE array or the Affymetrix Axiom^®^ UK Biobank array (12). Genotypes were imputed based on the Human Reference Consortium (HRC) panel, as described elsewhere (12). Variants with minor allele frequency (MAF) < 0.01 and imputation quality (info) scores < 0.3 were excluded from the analysis.

### SpiroMeta consortium

The SpiroMeta consortium meta-analysis comprised 75,696 individuals from 20 studies. Ten studies (N = 17,280) were imputed using 1000 Genomes Phase 1 reference panel (13, 14), nine studies (N = 37,919) were imputed using the Haplotype Reference Consortium (HRC) panel (12), and one study (N = 2077) was imputed using the HapMap CEU Build 36 Release 22. Two of the studies (ALSPAC and Raine) also provided data on children with an average age of 8 years (N = 5645). **Supplementary Tables 2** and **3** show the definitions of all abbreviations, study characteristics, details of genotyping platforms and imputation panels and methods. Measurements of spirometry for each study are as previously described (9, 15). Fourteen SpiroMeta studies had data on PEF so the sample size for replication of PEF signals was 51,555.

### The lung eQTL study

The lung expression quantitative trait loci (eQTL) study database has been described previously (16-18). It consists of non-tumour lung tissue samples from 1,111 individuals who had undergone lung resection surgery, mainly current or former smokers, genotyped on the Illumina Human1M-Duo BeadChip array.

### Statistical analysis

Spirometry-based lung function traits FEV_1_, FEV_1_/FVC, FVC, and PEF were pre-adjusted for age, age^2^, standing height (or sitting height in the sensitivity analysis) and smoking status and the residuals rank-transformed to normality using the rntransform function of the GenABEL package in R. To test each imputed variant for an interaction effect, a linear regression model with genotype (additive effect), sex, genotype-by-sex interaction, genotyping array and the first ten principal components included as covariates was implemented using Plink 2.0 software (https://www.cog-genomics.org/plink/2.0/). Genome-wide results were visualised using the R packages qqman, manhattanly and Circos v0.65. Region plots were generated using LocusZoom.

Following the genome-wide interaction study, sentinel SNPs were identified by selecting the SNP with the strongest sex interaction P value, excluding all SNPs (irrespective of LD) +/-1Mb, and then selecting the SNP with the strongest sex interaction P value from the remaining signals. This process was repeated until no SNPs with P<1 × 10^−6^ remained. Step-wise conditional analyses to identify independently associated variants within the 2Mb region surrounding the sentinel SNPs were then undertaken using GCTA software (19, 20). The SNPs with the strongest interaction P value at each independent locus were then selected for replication in SpiroMeta consortium studies.

Sentinel SNPs were also tested for association with lung function traits in males and females separately to provide sex-specific effect sizes.

Regression analysis to test genotype-by-sex interactions on height were conducted using a model including genotype (additive effect), age, age^2^, sex, genotyping array and the first ten principal components as covariates. Interactions between smoking status and genotype on lung function were tested using lung function traits transformed as described above (with sex included in the model instead of ever-smoking status). The linear regression model included genotype (additive effect), ever-smoking status, a genotype-by-smoking interaction term, genotyping array and the first ten principal components.

To test whether pubertal timing has differential effects on the association between SNPs and lung function in males and females, the regression model was adjusted for relative age at menarche in females and relative age at voice breaking in males. Relative age at voice breaking is categorised as earlier than average (1), around average (2) and later than average (3) in UK Biobank. Age at menarche is given as the participant’s age at menarche in years. To make these variables comparable, age at menarche was categorised as early (<12 years old), average (12-14 years old) and late (>14 years old) as in a previous study (21). As in the lung function analyses, ancestry-based principal components and genotyping array were included in all the regression models.

For the SpiroMeta consortium, summary statistics were generated by each contributing cohort separately according to the same analysis plan as the UK Biobank data. That is, the lung function traits were transformed as above and then tested for association with SNP-by-sex interaction terms, adjusting for sex and ancestry-based principal components. Meta-analysis of SpiroMeta cohorts was conducted using inverse-variance weighted fixed effects meta-analysis the metagen function of the meta package in R.

## Results

We tested 7,745,864 genome-wide variants with MAF ≥ 0.01 and imputation quality scores > 0.3 for genotype-by-sex interactions on lung function in 303,612 unrelated individuals of European ancestry from UK Biobank. Five independent signals were identified showing genome-wide significant (P<5 × 10^−8^) interaction with sex on at least one of four lung function traits (FEV_1_, FEV_1_/FVC, FVC, and PEF) with a further 21 SNPs showing suggestive significance (P<1 × 10^−6^) (**Table 1**, **Supplementary Figure 1**). The top three genome-wide significant signals had been previously reported for association with lung function: rs7697189 near the gene encoding hedgehog-interacting protein (*HHIP*) (interaction P = 3.15 × 10^−15^), rs9403386 near the gene encoding Adhesion G Protein-Coupled Receptor G6 (*ADGRG6*, previously known as *GPR126*) (interaction P = 4.56 × 10^−9^), and rs162185 downstream of the gene encoding transcription factor 21 (*TCF21*) (interaction P = 4.87 × 10^−9^) (22-27). Only rs355079 (interaction P = 8.84 × 10^−7^) showed an opposite direction of effect in males compared to females.

**Table 1.**
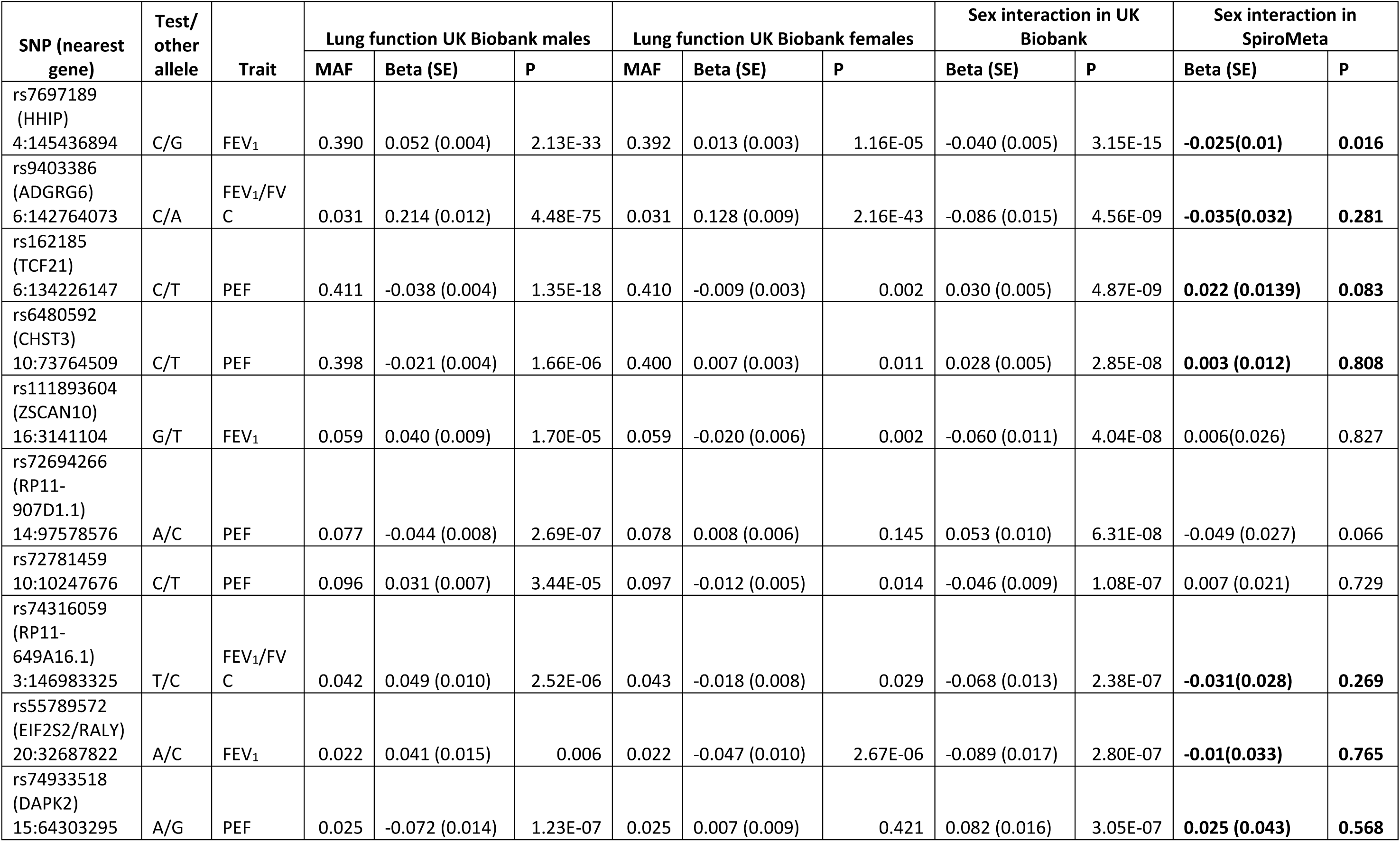
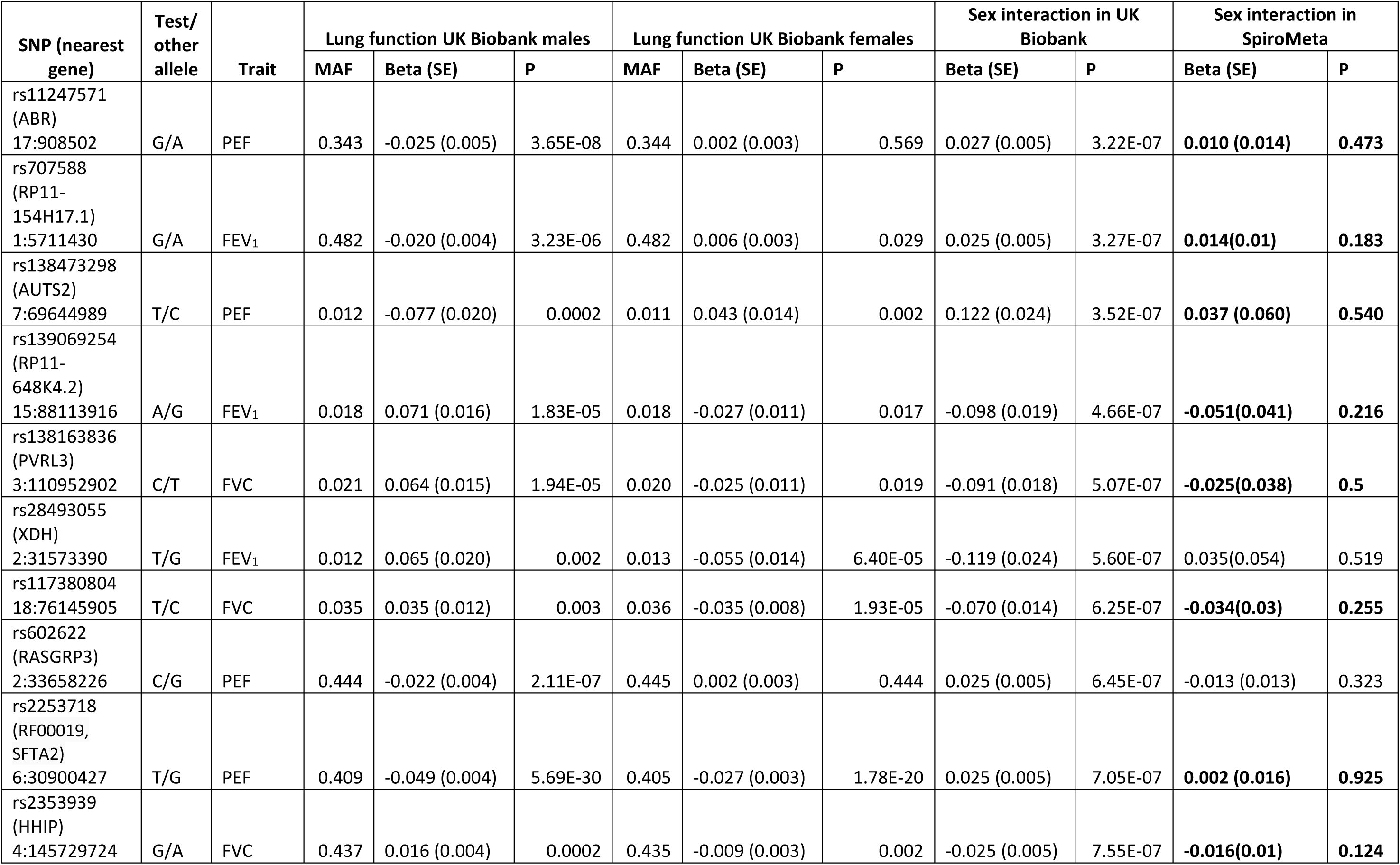
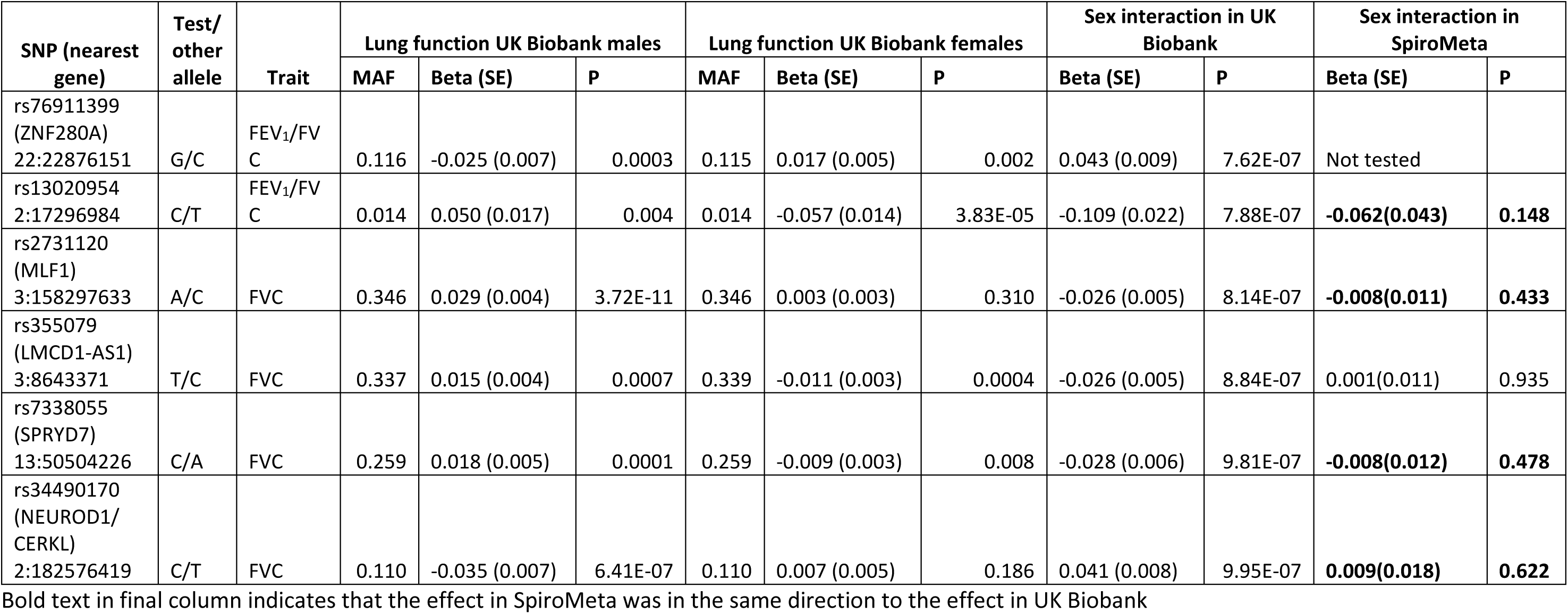
Association between sentinel SNPs and lung function in males and females, and genotype-by-sex interaction results on lung function in all

| SNP (nearest gene) | Test/<br>other allele | Trait | Lung function UK Biobank males |  |  | Lung function UK Biobank females |  |  | Sex interaction in UK Biobank |  | Sex interaction in SpiroMeta |  |
| --- | --- | --- | --- | --- | --- | --- | --- | --- | --- | --- | --- | --- |
|  |  |  | MAF | Beta (SE) | P | MAF | Beta (SE) | P | Beta (SE) | P | Beta (SE) | P |
| rs7697189<br>(HHIP)<br>4:145436894 | C/G | FEV<sub>1</sub> | 0.390 | 0.052 (0.004) | 2.13E-33 | 0.392 | 0.013 (0.003) | 1.16E-05 | -0.040 (0.005) | 3.15E-15 | **-0.025(0.01)** | **0.016** |
| rs9403386<br>(ADGRG6)<br>6:142764073 | C/A | FEV<sub>1</sub>/FV<br>C | 0.031 | 0.214 (0.012) | 4.48E-75 | 0.031 | 0.128 (0.009) | 2.16E-43 | -0.086 (0.015) | 4.56E-09 | **-0.035(0.032)** | **0.281** |
| rs162185<br>(TCF21)<br>6:134226147 | C/T | PEF | 0.411 | -0.038 (0.004) | 1.35E-18 | 0.410 | -0.009 (0.003) | 0.002 | 0.030 (0.005) | 4.87E-09 | **0.022 (0.0139)** | **0.083** |
| rs6480592<br>(CHST3)<br>10:73764509 | C/T | PEF | 0.398 | -0.021 (0.004) | 1.66E-06 | 0.400 | 0.007 (0.003) | 0.011 | 0.028 (0.005) | 2.85E-08 | **0.003 (0.012)** | **0.808** |
| rs111893604<br>(ZSCAN10)<br>16:3141104 | G/T | FEV<sub>1</sub> | 0.059 | 0.040 (0.009) | 1.70E-05 | 0.059 | -0.020 (0.006) | 0.002 | -0.060 (0.011) | 4.04E-08 | 0.006(0.026) | 0.827 |
| rs72694266<br>(RP11-907D1.1)<br>14:97578576 | A/C | PEF | 0.077 | -0.044 (0.008) | 2.69E-07 | 0.078 | 0.008 (0.006) | 0.145 | 0.053 (0.010) | 6.31E-08 | -0.049 (0.027) | 0.066 |
| rs72781459<br>10:10247676 | C/T | PEF | 0.096 | 0.031 (0.007) | 3.44E-05 | 0.097 | -0.012 (0.005) | 0.014 | -0.046 (0.009) | 1.08E-07 | 0.007 (0.021) | 0.729 |
| rs74316059<br>(RP11-649A16.1)<br>3:146983325 | T/C | FEV<sub>1</sub>/FV<br>C | 0.042 | 0.049 (0.010) | 2.52E-06 | 0.043 | -0.018 (0.008) | 0.029 | -0.068 (0.013) | 2.38E-07 | **-0.031(0.028)** | **0.269** |
| rs55789572<br>(EIF2S2/RALY)<br>20:32687822 | A/C | FEV<sub>1</sub> | 0.022 | 0.041 (0.015) | 0.006 | 0.022 | -0.047 (0.010) | 2.67E-06 | -0.089 (0.017) | 2.80E-07 | **-0.01(0.033)** | **0.765** |
| rs74933518<br>(DAPK2)<br>15:64303295 | A/G | PEF | 0.025 | -0.072 (0.014) | 1.23E-07 | 0.025 | 0.007 (0.009) | 0.421 | 0.082 (0.016) | 3.05E-07 | **0.025 (0.043)** | **0.568** |

| SNP (nearest gene) | Test/ other allele | Trait | Lung function UK Biobank males |  |  | Lung function UK Biobank females |  |  | Sex interaction in UK Biobank |  | Sex interaction in SpiroMeta |  |
| --- | --- | --- | --- | --- | --- | --- | --- | --- | --- | --- | --- | --- |
|  |  |  | MAF | Beta (SE) | P | MAF | Beta (SE) | P | Beta (SE) | P | Beta (SE) | P |
| rs11247571 (ABR)<br>17:908502 | G/A | PEF | 0.343 | -0.025 (0.005) | 3.65E-08 | 0.344 | 0.002 (0.003) | 0.569 | 0.027 (0.005) | 3.22E-07 | **0.010 (0.014)** | **0.473** |
| rs707588 (RP11-154H17.1)<br>1:5711430 | G/A | FEV<sub>1</sub> | 0.482 | -0.020 (0.004) | 3.23E-06 | 0.482 | 0.006 (0.003) | 0.029 | 0.025 (0.005) | 3.27E-07 | **0.014(0.01)** | **0.183** |
| rs138473298 (AUTS2)<br>7:69644989 | T/C | PEF | 0.012 | -0.077 (0.020) | 0.0002 | 0.011 | 0.043 (0.014) | 0.002 | 0.122 (0.024) | 3.52E-07 | **0.037 (0.060)** | **0.540** |
| rs139069254 (RP11-648K4.2)<br>15:88113916 | A/G | FEV<sub>1</sub> | 0.018 | 0.071 (0.016) | 1.83E-05 | 0.018 | -0.027 (0.011) | 0.017 | -0.098 (0.019) | 4.66E-07 | **-0.051(0.041)** | **0.216** |
| rs138163836 (PVRL3)<br>3:110952902 | C/T | FVC | 0.021 | 0.064 (0.015) | 1.94E-05 | 0.020 | -0.025 (0.011) | 0.019 | -0.091 (0.018) | 5.07E-07 | **-0.025(0.038)** | **0.5** |
| rs28493055 (XDH)<br>2:31573390 | T/G | FEV<sub>1</sub> | 0.012 | 0.065 (0.020) | 0.002 | 0.013 | -0.055 (0.014) | 6.40E-05 | -0.119 (0.024) | 5.60E-07 | 0.035(0.054) | 0.519 |
| rs117380804<br>18:76145905 | T/C | FVC | 0.035 | 0.035 (0.012) | 0.003 | 0.036 | -0.035 (0.008) | 1.93E-05 | -0.070 (0.014) | 6.25E-07 | **-0.034(0.03)** | **0.255** |
| rs602622 (RASGRP3)<br>2:33658226 | C/G | PEF | 0.444 | -0.022 (0.004) | 2.11E-07 | 0.445 | 0.002 (0.003) | 0.444 | 0.025 (0.005) | 6.45E-07 | -0.013 (0.013) | 0.323 |
| rs2253718 (RF00019, SFTA2)<br>6:30900427 | T/G | PEF | 0.409 | -0.049 (0.004) | 5.69E-30 | 0.405 | -0.027 (0.003) | 1.78E-20 | 0.025 (0.005) | 7.05E-07 | **0.002 (0.016)** | **0.925** |
| rs2353939 (HHIP)<br>4:145729724 | G/A | FVC | 0.437 | 0.016 (0.004) | 0.0002 | 0.435 | -0.009 (0.003) | 0.002 | -0.025 (0.005) | 7.55E-07 | **-0.016(0.01)** | **0.124** |

| SNP (nearest gene) | Test/other allele | Trait | Lung function UK Biobank males |  |  | Lung function UK Biobank females |  |  | Sex interaction in UK Biobank |  | Sex interaction in SpiroMeta |  |
| --- | --- | --- | --- | --- | --- | --- | --- | --- | --- | --- | --- | --- |
|  |  |  | MAF | Beta (SE) | P | MAF | Beta (SE) | P | Beta (SE) | P | Beta (SE) | P |
| rs76911399 (ZNF280A)<br>22:22876151 | G/C | FEV<sub>1</sub>/FVC | 0.116 | -0.025 (0.007) | 0.0003 | 0.115 | 0.017 (0.005) | 0.002 | 0.043 (0.009) | 7.62E-07 | Not tested |  |
| rs13020954<br>2:17296984 | C/T | FEV<sub>1</sub>/FVC | 0.014 | 0.050 (0.017) | 0.004 | 0.014 | -0.057 (0.014) | 3.83E-05 | -0.109 (0.022) | 7.88E-07 | **-0.062(0.043)** | **0.148** |
| rs2731120 (MLF1)<br>3:158297633 | A/C | FVC | 0.346 | 0.029 (0.004) | 3.72E-11 | 0.346 | 0.003 (0.003) | 0.310 | -0.026 (0.005) | 8.14E-07 | **-0.008(0.011)** | **0.433** |
| rs355079 (LMCD1-AS1)<br>3:8643371 | T/C | FVC | 0.337 | 0.015 (0.004) | 0.0007 | 0.339 | -0.011 (0.003) | 0.0004 | -0.026 (0.005) | 8.84E-07 | 0.001(0.011) | 0.935 |
| rs7338055 (SPRYD7)<br>13:50504226 | C/A | FVC | 0.259 | 0.018 (0.005) | 0.0001 | 0.259 | -0.009 (0.003) | 0.008 | -0.028 (0.006) | 9.81E-07 | **-0.008(0.012)** | **0.478** |
| rs34490170 (NEUROD1/CERKL)<br>2:182576419 | C/T | FVC | 0.110 | -0.035 (0.007) | 6.41E-07 | 0.110 | 0.007 (0.005) | 0.186 | 0.041 (0.008) | 9.95E-07 | **0.009(0.018)** | **0.622** |

Bold text in final column indicates that the effect in SpiroMeta was in the same direction to the effect in UK Biobank

**Figure 1.**
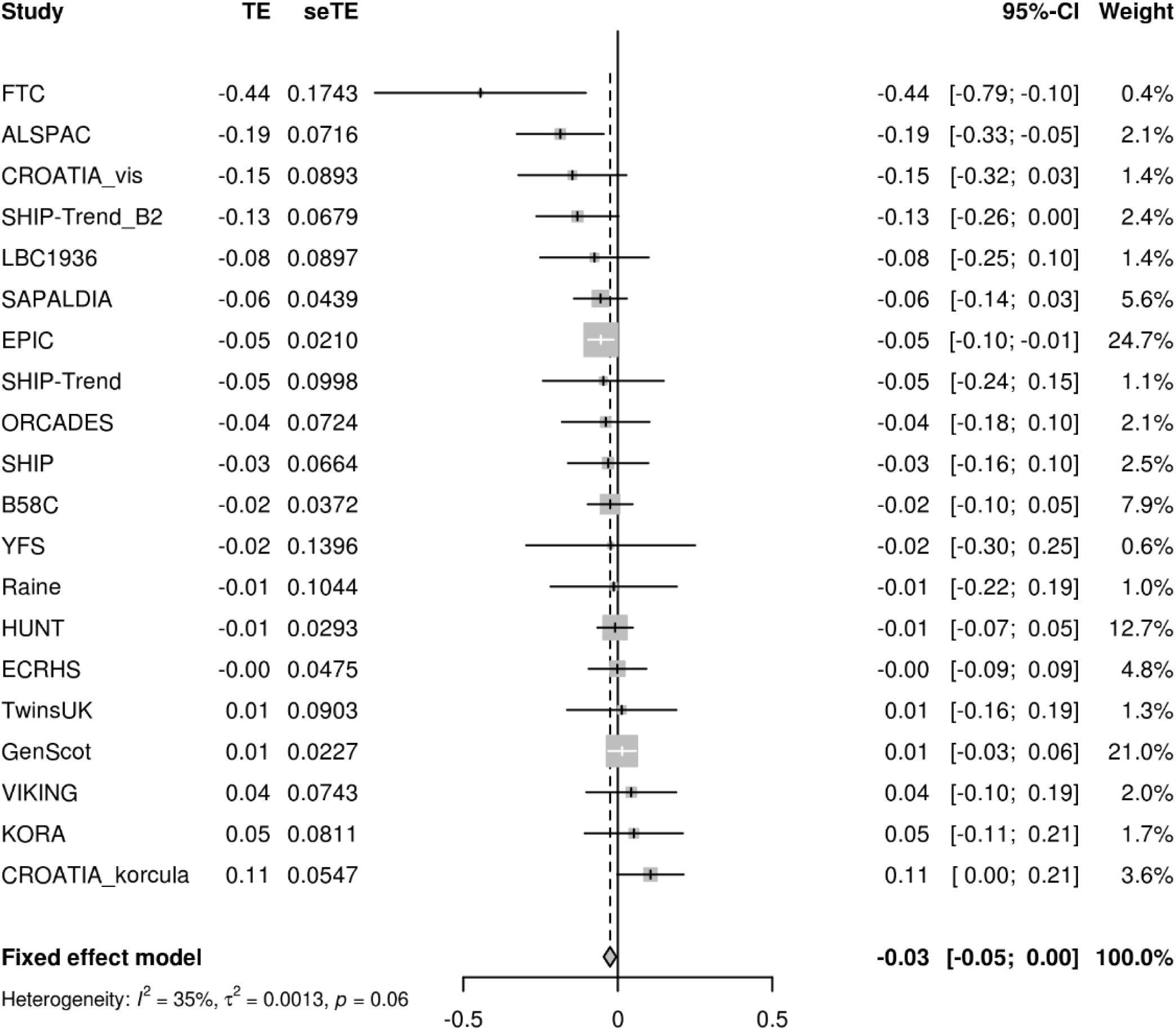
Forest plot showing the beta-coefficients (test effects, TE) and standard errors for the interaction between rs7697189 and sex on FEV_1_ in 20 cohorts of the SpiroMeta consortium. The overall effect size from fixed effects meta-analysis is represented by the diamond.

We sought evidence for replication of all 26 signals in up to 75,696 individuals from 20 cohorts of the SpiroMeta consortium. One variant, rs76911399, was poorly imputed in SpiroMeta cohorts (effective sample size, N_eff_ = 41,135) and was excluded. Of the UK Biobank genotype-by-sex interaction signals, one SNP showed a nominally significant (P < 0.05) interaction with sex on lung function in SpiroMeta cohorts: rs7697189 (near *HHIP*) (replication interaction P = 0.016) (**Table 1**, **Figure 1**). A further 18 signals were not significant in SpiroMeta but exhibited the same direction of interaction effect as in UK Biobank.

The strongest sex interaction signal came from rs7697189 on FEV_1_ (P = 3.15 × 10^−15^ in UK Biobank and P=0.016 in SpiroMeta). The minor (C) allele of rs7697189 has a larger effect on lung function in males (β = 0.052 [SE 0.004], P = 2.13 × 10^−33^) compared to females (β = 0.013 [SE 0.003], P = 1.16 × 10^−5^) (**Table 1)**. This SNP resides upstream of the *HHIP* gene and is in linkage disequilibrium with two previously reported lung function-associated sentinel SNPs, rs13141641 (27, 28) (r^2^ = 0.91) and rs13116999 (28) (r^2^ = 0.56). SNP rs7697189 was also genome-wide and suggestively significant for interactions with sex on PEF and FEV_1_/FVC respectively, but did not meet the threshold for suggestive significance in FVC (P = 8.71 × 10^−5^) (**Supplementary Table 4**, **Figure 2**).

**Figure 2.**
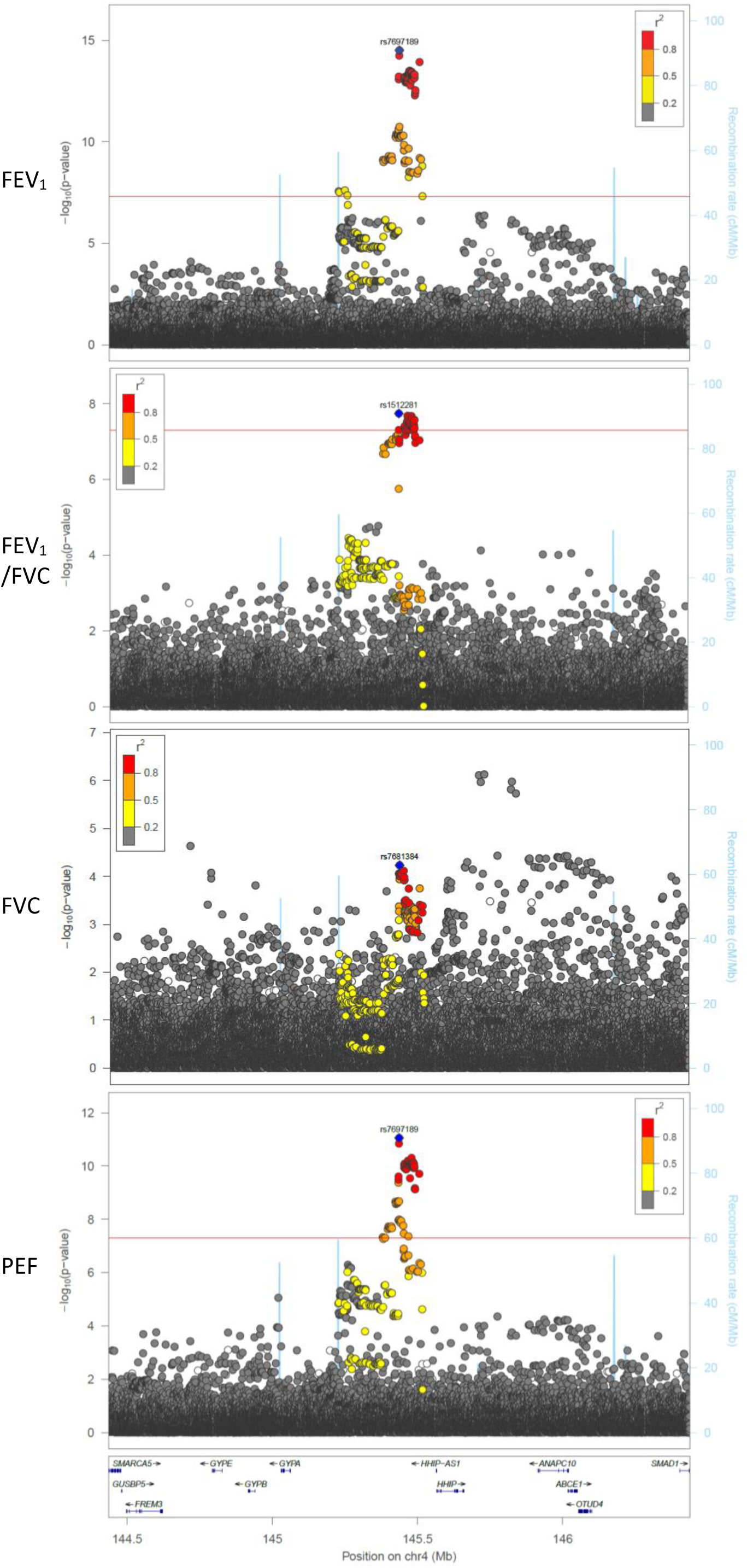
Region plots showing genotype-by-sex interaction results within the *HHIP* region for lung function traits (FEV_1_, FEV_1_/FVC, FVC, and PEF). The SNP with the strongest association in the rs7697189-proximal region is represented by a blue diamond. The FEV_1_ and PEF sentinels are rs7697189, the FEV_1_/FVC sentinel is rs1512281 (R^2^ = 0.95 with rs7697189), and the FVC sentinel is rs7681384 (R^2^ = 0.57 with rs7697189). Note that there is an independent suggestively significant signal from rs2353939 and surrounding SNPs for FVC, but this did not replicate in SpiroMeta cohorts. All other SNVs are colour coded according to their linkage disequilibrium (R^2^) with the sentinel SNP (as shown in the key). All imputed SNVs are plotted irrespective of MAF, demonstrating that rarer variants are not exhibiting significant interactions with sex on lung function. The locations of genes in the region are shown in the lower panel of each plot. Recombination rate is represented by the blue lines.

### rs7697189 interacts with sex on lung function independently of height, smoking and pubertal timing

As SNPs in *HHIP* are also reported to be associated with height (29) and increased height is associated with increased lung function, it is possible that rs7697189 has differential effects on lung function in males and females through differential effects on height. However, the association of rs7697189 with standing height was not modified by sex in a combined analysis of UK Biobank males and females with a genotype-by-sex interaction term (interaction P = 0.806). We also conducted a sensitivity analysis showing that the rs7697189-by-sex interaction on FEV_1_ remained significant after adjustment for sitting height (β = −40.04 [SE = 0.005], P = 1.97 × 10^−;15^).

Amongst the 303,612 UK Biobank participants in this study, the proportion of ever-smokers was higher in males (52.8%) than females (40.3%) (**Supplementary Table 1**). A larger effect of rs7697189 on lung function in males compared to females could arise if there was an interaction effect with smoking. However, there was no interaction between rs7697189 and ever-smoking status on FEV_1_ in this study (interaction P = 0.63).

SNP rs7697189, and correlated SNPs in the region, have been shown to be associated with expression levels of Hedgehog-interacting protein (*HHIP*) in lung tissue (30). HHIP is a critical protein during early development and *HHIP* variants have been associated with lung function in infancy (31). We tested whether *HHIP* SNPs also have differential effects on lung function in females compared to males in childhood using data from children with an average age of 8 years in the ALSPAC and Raine studies (N = 5645). No significant association between rs7697189 and FEV_1_, and no significant interaction between rs7697189 and sex on FEV_1_ was detected, possibly due to the much smaller sample size of the childhood cohorts (**Supplementary Figure 2**). Finally, as pubertal timing has been associated with adult lung function (15), we tested for an effect of relative age at puberty on the association between rs7697189 and lung function in a sex-stratified analysis. The association between *HHIP* SNPs and lung function was adjusted for relative age at voice breaking in males and for age at menarche in females, but neither adjustment changed the effect estimate of the SNPs on lung function (**Supplementary Table 5).**

### rs7697189 is associated with HHIP expression, but no interaction with sex

It is possible that rs7697189 interacts with sex on lung function through differential effects on *HHIP* expression. We confirm that rs7697189 is associated with *HHIP* expression in lung tissue but we do not detect an interaction with sex on *HHIP* expression (**Supplementary Table 6**). However, *HHIP* (in all samples irrespective of genotype at rs7697189) does show differential expression between males and females, with females showing higher expression (**Supplementary Table 7**).

## Discussion

We identified a genome-wide significant genotype-by-sex interaction signal at a locus previously reported for association with lung function upstream of the *HHIP* gene (rs7697189, FEV_1_ interaction P = 3.15 × 10^−;15^). The signal was nominally significant in 75,696 individuals from 20 independent studies of the SpiroMeta consortium. We demonstrated that the differential effects of this SNP in males and females (untransformed FEV_1_ β = 0.028 [SE 0.0022] litres in males vs β = 0.009 [SE 0.0014] litres in females) was not mediated by effects on height, smoking behaviour or pubertal age.

SNPs at the *HHIP* locus also exhibited genome-wide or suggestive significant interactions with sex on two additional lung function traits in UK Biobank: FEV_1_/FVC and PEF (P = 8.98 × 10^−;8^ and P = 8.78 × 10^−;12^ respectively). Stratified analyses in males and females separately demonstrated that these SNPs appeared to have a stronger effect on lung function in males compared to females. There was no interaction between these SNPs and ever-smoking status on lung function in UK Biobank, suggesting that the stronger effect in males is not due to differences in smoking behaviour. We also demonstrate that an association between these SNPs and height is not modified by sex, suggesting that differential effects on height in males and females do not explain the genotype-by-sex interaction on lung function.

The genome-wide significant sex interaction locus is located upstream of the *HHIP* gene, a region previously reported to be associated with lung function (23, 26) and *HHIP* gene expression (30). The *HHIP* gene encodes hedgehog-interacting protein, a negative regulator of hedgehog signalling. The hedgehog signalling pathway regulates numerous physiological processes such as growth, self-renewal, cell survival, differentiation, migration, and tissue polarity and plays a vital role in the morphogenesis of lung and other organs (32). Hedgehog signalling has also been shown to participate in regulation of stem and progenitor cell populations in adult tissues, impacting tissue homeostasis and repair (33). SNP rs7697189, showing the strongest sex interaction on lung function in our study, is in strong linkage disequilibrium (R^2^ = 0.93) with SNPs residing in an *HHIP* enhancer region (30). These enhancer-region SNPs also exhibit genome-wide significant genotype-by-sex interactions on lung function in our data. According to previous reports, the alleles associated with decreased lung function are also associated with reduced enhancer activity and reduced *HHIP* expression in lung tissues, providing a potential mechanism by which SNPs in this linkage disequilibrium block might modulate hedgehog signalling in the lungs (30). However, in our study, though *HHIP* was expressed at lower levels in males compared to females in lung tissue, the association between rs7697189 and *HHIP* expression was not modified by sex. This may be because there is no sex differential effect on expression, or the study might have been underpowered to detect an interaction effect (based on 472 males and 566 females). It is therefore still not clear why SNPs upstream of *HHIP* would be showing different effects in males and females.

Investigating the effects of HHIP at different stages of development by sex may help to shed light on this conundrum. In our study we had access to genetic and lung function data from 5645 children with an average age of 8 years. Though underpowered to detect the association between rs7697189 and FEV_1_ seen in UK Biobank adults, the lack of a similar trend in children suggests that *HHIP* variants may have differential effects at different developmental stages (though the genotype-by-sex interaction is in the same direction as in adults). We also looked for an effect of timing of puberty on the association between rs7697189 and lung function in adults, but adjustment for relative age of voice breaking in males and relative age at menarche in females made no difference to the relationship between rs7697189 and lung function.

We identified four additional genome-wide significant (interaction P<5×10^−;8^) sex-by-genotype interactions on lung function in our discovery analysis in UK Biobank, with a further 21 that met a less stringent threshold of interaction (P<1×10^−;6^). As far as we are aware, this is the first genome-wide sex-by-genotype interaction study for lung function traits. We did not find a significant genotype-by-sex interaction on lung function or COPD at the *CELSR1* locus (interaction P = 0.525 and P = 0.503 respectively) previously reported to have sex-specific effects on risk of COPD (10).

In conclusion, we have identified a novel genotype-by-sex interaction at SNPs at a putative enhancer region upstream of the hedgehog-interacting protein (*HHIP*) gene. Establishing the mechanism by which *HHIP* has sex differential effects on lung function will be important for our understanding of COPD and for realising the potential of precision medicine by optimising treatment in males and females.

## Study contributions

### Contributed to the conception and design of the study

M.O.,C.J.,M.I.,P.H.,C.L.,Y.B.,D.D.S.,K.H.,R.E.,B.S.,O.P.,J.M.S.,I.R.,K.Pietiläinen,M.K.,O.T.R,G.L.H.,P.D.S., C.E.P.,J.K.,T.L.,I.J.D.,J.F.W.,N.Probst-Hensch,N.W.,H.Z.,D.P.S.,B.M.B.,C.H.,I.P.H.,M.D.T.,L.V.W.

### Undertook data analysis

K.F.,M.O.,C.A.M.,A.L.G.,J.L.,A.R.,M.R.M.,R.G.,S.W.,M.I.,S.M.,T.S.B.,L.P.,C.F.,S.E.H,C.A.W.,L.L.,T.P.,R.E.F.,D.K.,M.V.,N.P.,K.P.,N.Probst-Hensch,D.P.S.,B.M.B.,M.D.T.,L.V.W.

### Contributed to data acquisition and/or interpretation

N.S.,J.L.,R.G.,S.W.,M.I.,P.H.,S.E.H.,C.A.W.,L.L.,R.E.F.,D.K.,C.M.,C.L.,D.E.,S.R.,A.L.,K.Hveem,C.W.,R.E.,B. S.,L.K.,P.K.J.,M.M.,O.P.,J.M.S.,I.R.,T.R.,K.Pietiläinen,M.K.,O.T.R.,G.L.H.,P.D.S.,C.E.P.,J.K.,T.L.,V.V.,I.J.D., D.J.,J.F.W.,T.S.,N.Probst-Hensch,N.W.,H.Z.,J.H.,D.P.S.,B.M.B.,C.H.,I.P.H.,M.D.T.,L.V.W.

### Drafted the manuscript

K.F.,C.M.,L.V.W

## Funding and acknowledgements

I.P.H.: The research was partially supported by the NIHR Nottingham Biomedical Research Centre; the views expressed are those of the author(s) and not necessarily those of the NHS, the NIHR or the Department of Health.

M.D.T.: M.D. Tobin is supported by a Wellcome Trust Investigator Award (WT202849/Z/16/Z). M.D. Tobin and L.V. Wain have been supported by the MRC (MR/N011317/1). The research was partially supported by the NIHR Leicester Biomedical Research Centre; the views expressed are those of the author(s) and not necessarily those of the NHS, the NIHR or the Department of Health.

L.V.W.: L.V. Wain holds a GSK/British Lung Foundation Chair in Respiratory Research.

### Funding and acknowledgment statements from individual cohorts

The UK Biobank analysis was conducted under approved UK Biobank data application number 648.

CROATIA-Korcula/Split/Vis: MRC, University Unit Programme Grant (MC_PC_U127592696), European Union, EUROSPAN project (contract no. LSHG-CT-2006-018947), Croatian Ministry of Science (216-1080315-0302) (I.R.)

FTC: Academy of Finland (308248, 312073) (J.K.), Sigrid Juselius Foundation (J.K.), Academy of Finland (213506) (T.R.), Academy of Finland (272376, 314383, 266286) (K.P.), Finnish Medical Foundation (K.P.), Novo Nordisk Foundation (K.P.), Finnish Diabetes Research Foundation (K.P.), State Research Funds (K.P.), University of Helsinki (K.P.)

Generation Scotland: MRC, University Unit Programme Grant (MC_PC_U127592696), Wellcome Trust, Strategic Award 104036/Z/14/Z, Chief Scientist Office (CZD/16/6), Scottish Funding Council (HR03006)

HUNT: Stiftelsen Kristian Gerhard Jebsen (K.H.), The Liaison Committee for education, research and innovation in Central Norway (K.H., B.M.B.), NIH (HL135824, HL109946, HL127564) (C.W.): The Nord-Trøndelag Health Study (The HUNT Study) is a collaboration between HUNT Research Center (Faculty of Medicine and Health Sciences, NTNU, Norwegian University of Science and Technology), Nord-Trøndelag County Council, Central Norway Regional Health Authority, and the Norwegian Institute of Public Health.

KORA: MC-Health, LMUinnovativ: The KORA study was initiated and financed by the Helmholtz Zentrum München – German Research Center for Environmental Health, which is funded by the German Federal Ministry of Education and Research (BMBF) and by the State of Bavaria. Furthermore, KORA research was supported within the Munich Center of Health Sciences (MC-Health), Ludwig-Maximilians-Universität, as part of LMUinnovativ.

LBC1936: Biotechnology and Biological Sciences Research Council (BBSRC) (BB/F019394/1), Age UK (The Disconnected Mind Project) (DCM and DCM PHASE 2), Cross Council Lifelong Health and Wellbeing Initiative (MR/K026992/1): We gratefully acknowledge the contribution of co-author Professor John M. Starr, who died prior to the publication of this manuscript.

ORCADES/VIKING: Chief Scientist Office (CZB/4/276, CZB/4/710) (J.F.W.), MRC (MC_UU_00007/10) (J.F.W.), MRC (MR/N013166/1) (S.M.), EU FP6 (LSHG-CT-2006-018947) (J.F.W.), Royal Society (URF to J.F.W.): The Viking Health Study – Shetland (VIKING) DNA extractions and genotyping were performed at the Edinburgh Clinical Research Facility, University of Edinburgh. We would like to acknowledge the invaluable contributions of the research nurses in Shetland, the administrative team in Edinburgh and the people of Shetland. The Orkney Complex Disease Study (ORCADES) DNA extractions were performed at the Wellcome Trust Clinical Research Facility in Edinburgh. We would like to acknowledge the invaluable contributions of the research nurses in Orkney, the administrative team in Edinburgh and the people of Orkney.

Raine: National Health and Medical Research Council of Australia (NHMRC) (572613, 403981, 003209), Canadian Institutes of Health Research (CIHR) (MOP-82893), Raine Medical Research Foundation: The Raine study would like to acknowledge the continued contribution of Raine Study participants and their families, Raine Study research staff for cohort coordination and data collection, NHMRC for long term funding over last 29 years, the Raine Medical Research Foundation, UWA Faculty of Medicine, Dentistry and Health Sciences, Telethon Kids Institute, Women’s and Infants’ Research Foundation, Curtin University, Edith Cowan University, Murdoch University, and University of Notre Dame for providing funding for Core Management of the Raine Study, the Western Australian DNA Bank (National Health and Medical Research Council of Australia National Enabling Facility) for providing assistance in handling and storing the Raine Study samples, and Pawsey Supercomputing Centre with funding from Australian Government and the Government of Western Australia for providing computation resource to carry out analyses required.

SAPALDIA: Swiss National Science Foundation (33CS30-148470/1&2, 33CSCO-134276/1, 33CSCO-108796, 324730_135673, 3247BO-104283, 3247BO-104288, 3247BO-104284, 3247-065896, 3100-059302, 3200-052720, 3200-042532, 4026-028099, PMPDP3_129021/1, PMPDP3_141671/1) (SAPALDIA1 to SAPALIDA5), Canton’s government of Aargau, Basel-Stadt, Basel-Land, Geneva, Luzern, Ticino, Valais, and Zürich, Federal Offices of Environment, of Public Health, and of Roads and Transport, Cantonal lung leages of Basel Stadt/ Basel Landschaft, Geneva, Ticino, Valais, Graubünden and Zurich, Horizon 2020 (633212) (ALEC Project), European Commission (308610, 018996) (GABRIEL Project), Freie Akademaische Gesellschaft (N.P.), UBS Wealth Foundation (N.P.), Wellcome Trust (WT 084703MA) (GABRIEL Project), Talecris Biotherapeutics GmbH (N.P.), Abbott Diagnostics (N.P.)

SHIP/SHIP_Trend/SHIP_Trend_B2: German Federal Ministry of Education and Research (01ZZ9603, 01ZZ0103, and 01ZZ0403), German Research Foundation (GR 1912/5-1)

YFS: Academy of Finland (286284, 134309 (Eye), 126925, 121584, 124282, 129378 (Salve), 117787 (Gendi), and 41071 (Skidi)), Social Insurance Institution of Finland, Competitive State Research Financing of the Expert Responsibility area of Kuopio, Tampere and Turku University Hospitals, Juho Vainio Foundation, Paavo Nurmi Foundation, Finnish Foundation for Cardiovascular Research, Finnish Cultural Foundation, The Sigrid Juselius Foundation, Tampere Tuberculosis Foundation, Emil Aaltonen Foundation, Yrjö Jahnsson Foundation, Signe and Ane Gyllenberg Foundation, Diabetes Research Foundation of Finnish Diabetes Association, EU Horizon 2020 (755320 for TAXINOMISIS), European Research Council (742927 for MULTIEPIGEN), Tampere University Hospital Supporting Foundation

## Competing interests

### The following authors report potential competing interests

L.V.W.: Louise V. Wain has received grant support from GSK.

M.D.T.: Martin D. Tobin has received grant support from GSK.

I.P.H.: Ian P. Hall has received support from GSK and BI.

