## Supplementary Figures and Tables for "SNPs associated with *HHIP* expression have differential effects on lung function in males and females"

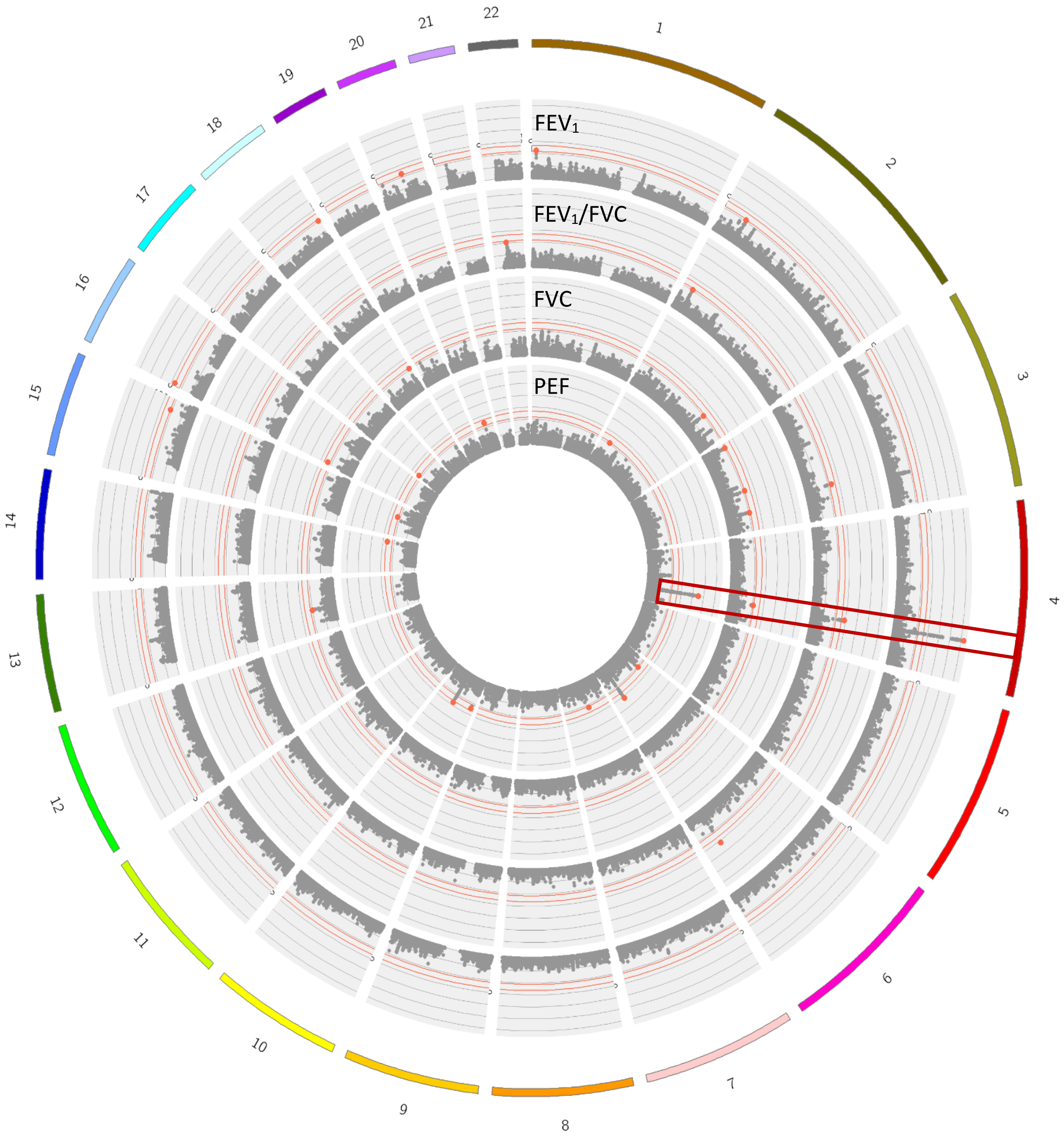

**Supplementary Figure 1**. A Circos plot showing genome-wide interaction results between imputed SNPs (MAF>0.01) and sex on four lung function traits ($FEV_1$, $FEV_1$/FVC, FVC and PEF) in 303,612 UK Biobank participants. Sentinel SNPs exhibiting genome-wide or suggestive significant interaction with sex on lung function (P < 1e-6) are highlighted as red dots. The *HHIP* locus signal is additionally highlighted by the red box. The red lines show genome-wide and suggestive significance boundaries. The Y axes represent $-\log_{10}$ P values ranging between 2 and 15.

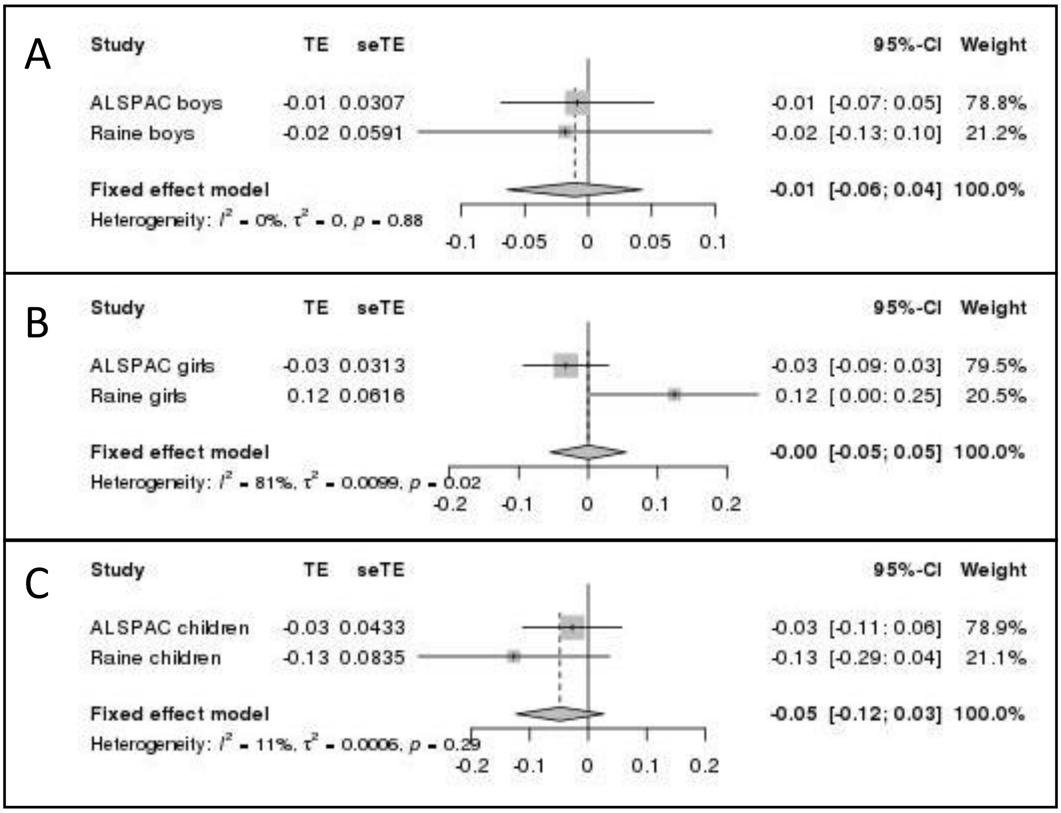

**Supplementary Figure 2.** Forest plots showing the beta-coefficients (test effects, TE) and standard errors for the association between rs7697189 and $FEV_1$ in boys (A) and girls (B) with an average age of 8 years old from the ALSPAC and Raine cohorts, and for the genotype-by-sex interaction on $FEV_1$ (C). The outcome of fixed effects meta-analyses are represented by the diamonds. The coded allele is always the C allele and the MAF in ALSPAC boys and girls is 0.39 and in Raine boys and Raine girls is 0.41.

**Supplementary Table 1: UK Biobank demographics**

| | All | Females | Males |
|---|---|---|---|
| **N Total** | 303,612 | 168,137 | 135,475 |
| **Age range (y) at lung function measurement** | 39-72 | 39-71 | 39-72 |
| **Mean age, y (s.d.)** | 56.47 (7.98) | 56.26 (7.88) | 56.73 (8.09) |
| **Mean height, cm (s.d.)** | 168.61 (9.13) | 162.77 (6.18) | 175.84 (6.70) |
| **Mean $FEV_1$, L (s.d.)** | 2.84 (0.76) | 2.44 (0.51) | 3.34 (0.72) |
| **Mean FVC, L (s.d.)** | 3.74 (0.96) | 3.18 (0.62) | 4.43 (0.86) |
| **Mean $FEV_1$/FVC (s.d.)** | 0.76 (0.06) | 0.77 (0.06) | 0.75 (0.07) |
| **Mean PEF, L/min (s.d.)** | 406.72 (117.43) | 342.37 (74.69) | 486.58 (111.63) |
| **N never smokers** | 164,327 | 100,375 | 63,952 |
| **N ever smokers** | 139,285 | 67,762 | 71,523 |
| **UK BiLEVE array** | 44,459 | 21,992 | 22,467 |
| **UK Biobank array** | 259,153 | 146,145 | 113,008 |

**Supplementary Table 2: SpiroMeta Studies**

ALSPAC (Avon Longitudinal Study of Parents and Children); B58C (B58C-T1DGC, British 1958 Birth Cohort–Type 1 Diabetes Genetics Consortium; B58C-GABRIEL British 1958 Birth Cohort–GABRIEL consortium; B58C-WTCCC, British 1958 Birth Cohort–Wellcome Trust Case Control Consortium); the CROATIA-Korcula study; the CROATIA-Vis study; ECRHS (European Community Respiratory Health Survey); EPIC population based, European Prospective Investigation into Cancer and Nutrition Cohort; FinnTwin (Finnish Twin Cohort); GS:SFHS, Generation Scotland: Scottish Family Health Study; HUNT (Nord-Trøndelag Health Study); KORA, Cooperative Health Research in the Region of Augsburg; LBC1936, Lothian Birth Cohort 1936; ORCADES, Orkney Complex Disease Study; Raine (Western Australian Pregnancy Cohort); SHIP, Study of Health in Pomerania; SHIP-TREND; SHIP-TREND-B2 (SHIP-TREND batch 2); TwinsUK; VIKING; YFS, the Young Finish Study. The total size in this table is not exactly equal to the maximum sample size given in the main text, since some studies had subtly different subsets of individuals entering each of the four lung function trait GWAS.

| Study name | N Total | N male | N female | Age range (y) at lung function measurement | Mean age, y (s.d.) | Mean height, cm (s.d.) | Mean $FEV_1$, L (s.d.) | Mean FVC, L (s.d.) | Mean $FEV_1$/FVC (s.d.) | Mean PEF, L/sec (s.d.) | N never smokers | N ever smokers | Genotyping Platform |
|---|---|---|---|---|---|---|---|---|---|---|---|---|---|
| ALSPAC | 1707 | 607 | 1100 | 22-26 | 23.98 (0.80) | 171.32 (9.37) | 3.85 (0.82) | 4.52 (1.03) | 0.86 (0.06) | -- | 634 | 1073 | Illumina human660W quad chip |
| ALSPAC children | 4426 | 2240 | 2186 | 7-10 | 8.65 (0.25) | 132.41 (5.70) | 1.69 (0.26) | 1.93 (0.32) | 0.88 (0.07) | -- | 2610* | 1816* | Illumina human660W quad chip |
| B58C | 5934 | 2955 | 2979 | 44-45 | 45.12 (0.38) | 169.43 (9.29) | 3.30 (0.76) | 4.19 (0.98) | 0.79 (0.08) | -- | 1709 | 4225 | Illumina 550k/610k |
| CROATIA-Korcula | 2577 | 944 | 1633 | 18-98 | 53.51 (15.44) | 169.3 (9.38) | 2.89 (0.15) | 3.35 (0.98) | 0.86 (0.15) | 370.1 (136.74) in L/min | 1275 | 1302 | Illumina HumanHap370CNV duo chip |
| CROATIA-Vis | 925 | 390 | 535 | 18-88 | 55.90 (15.51) | 167.80 (9.88) | 3.42 (1.21) | 4.41 (1.42) | 0.77 (0.09) | 6.33 (2.89) | 388 | 537 | Illumina Infinium HumanHap300 BeadChip |
| ECRHS | 2077 | 991 | 1086 | 19-48 | 33.93 (7.19) | 170.36 (9.6) | 3.68 (0.84) | 4.54 (1.05) | 0.81 (0.08) | -- | 909 | 1167 | Illumina 610k |
| EPIC population based | 18662 | 8760 | 9902 | 39-79 | 59.13 (9.2) | 167.2 (9.1) | 2.51 (0.74) | 3.06 (0.92) | 0.83 (0.11) | 3.65 (1.23) | 9532 | 11239 | Affymetrix UKBioBank Axiom |
| FinnTwin | 451 | 88 | 363 | 22-61 | 43.46 (15.53) | 166 (9.62) | 2.86 (1.08) | 3.55 (1.25) | 0.80 (0.08) | -- | 147 | 304 | Batch1: Illumina Human610-Quad v1.0 B, Human670-QuadCustom v1.0 A & Batch2: Illumina HumanCoreExome (12 v1.0 B, 12 v1.1 A, 24 v1.0 A, 24 v1.1 A) |
| GS:SFHS | 15867 | 6570 | 9297 | 18-93 | 46.84 (14.58) | 168.4 (9.43) | 2.97 (0.04) | 3.88 (1.00) | 0.76 (0.04) | -- | 8571 | 7296 | Illumina OmniExpress+Exome |
| HUNT | 9095 | 4205 | 4890 | 19-98 | 50.80 (16.09) | 170.58 (9.16) | 3.19 (1.01) | 4.13 (1.17) | 0.77 (0.09) | 7.40 (2.51) | 3360 | 5735 | Illumina (HumanCoreExome12 v1.0, HumanCoreExome12 v1.1 and UM HUNT Biobank v1.0) |
| KORA | 1234 | 573 | 661 | 41-62 | 51.71 (5.75) | 169.71 (9.28) | 3.34 (0.82) | 4.31 (1.01) | 0.78 (0.06) | 7.4 (2.08) | 445 | 789 | Affymetrix Axiom |

| Study name | N Total | N male | N female | Age range (y) at lung function measurement | Mean age, y (s.d.) | Mean height, cm (s.d.) | Mean $FEV_1$, L (s.d.) | Mean FVC, L (s.d.) | Mean $FEV_1$/FVC (s.d.) | Mean PEF, L/sec (s.d.) | N never smokers | N ever smokers | Genotyping Platform |
|---|---|---|---|---|---|---|---|---|---|---|---|---|---|
| LBC1936 | 1002 | 509 | 493 | 68-71 | 69.55 (0.84) | 166.48 (8.93) | 2.37 (0.69) | 3.05 (0.87) | 0.78 (0.10) | 352.73 (133.7) in L/min | 466 | 536 | Illumina 610-Quadv1 |
| ORCADES | 2215 | 870 | 1344 | 16-100 | 54.09 (15.29) | 167 (9.00) | 2.84 (0.85) | 3.55 (0.99) | 0.80 (0.08) | 409.54 (127.73) in L/min | 1215 | 750 | Illumina Hap300, Illumina Omni1 & Illumina OmniX |
| Raine | 789 | 391 | 398 | 20-25 | 22.2 (0.74) | 173 (10.0) | 3.98 (0.82) | 4.8 (1.06) | 0.83 (0.06) | -- | 637 | 152 | Illumina Human660W-Quad BeadChip |
| Raine children | 1219 | 630 | 589 | 7-9 | 8.09 (0.34) | 129 (6.00) | 1.56 (0.25) | 1.65 (0.28) | 0.95 (0.05) | -- | -- | -- | Illumina Human660W-Quad BeadChip |
| SAPALDIA | 3981 | 2008 | 1973 | 29-73 | 51.30 (11.09) | 169.49 (9.11) | 3.24 (0.83) | 4.33 (1.04) | 0.75 (0.08) | 7.77 (2.24) | 1843 | 2138 | Illumina 610k quad, Illumina OmniExpressExome |
| SHIP | 1752 | 853 | 899 | 20-80 | 47.10 (13.63) | 169.7 (9.14) | 3.29 (0.89) | 3.88 (1.03) | 0.85 (0.06) | 7.31 (2.07) | 818 | 934 | Affymetrix SNP 6.0 |
| SHIP-TREND | 800 | 360 | 440 | 21-81 | 51.22 (13.34) | 169.9 (9.02) | 3.30 (0.87) | 4.14 (1.06) | 0.80 (0.06) | 6.57 (2.09) | 341 | 459 | Illumina Human Omni 2.5 |
| SHIP-TREND-B2 | 1707 | 894 | 813 | 20-81 | 53.07 (14.65) | 170.4 (9.34) | 3.25 (0.92) | 4.12 (1.12) | 0.79 (0.06) | 6.64 (2.13) | 622 | 1085 | Illumina CoreExome v1.0 |
| TwinsUK | 4227 | 380 | 3847 | 16-81 | 47.8 (12.46) | 163.69 (7.22) | 2.86 (0.67) | 3.54 (0.75) | ? | -- | 1806 | 2419 | Illumina arrays (HumanHap300, HumanHap610Q, 1M-Duo and 1.2MDuo 1M) |
| VIKING | 2181 | 842 | 1262 | 18-93 | 49.93 (15.26) | 168 (9.00) | 3.07 (0.82) | 4.03 (0.98) | 0.76 (0.10) | 449.28 (131.43) in L/min | 1153 | 931 | Illumina OmniExpress Exome |
| YFS | 419 | 198 | 221 | 30-47 | 38.88 (5.07) | 172.25 (8.90) | 3.73 (0.75) | 4.68 (0.99) | 0.8 (0.06) | -- | 233 | 186 | Illumina 670k custom |

*Exposure to cigarette smoke from birth to 8 years old

### Supplementary Table 3: SpiroMeta analysis method

| Study name | Individual call rate filter (applied before imp'n) | SNP call rate filter (applied before imp'n) | SNP HWE *P* filter (applied before imp'n) | SNP MAF filter (applied before imp'n) | Other filters | No of SNPs after filtering (before imp'n) | Imputation software and version | Reference panel used for imp'n |
|---|---|---|---|---|---|---|---|---|
| ALSPAC | 97% | 95% | 1.00E-06 | 0.01 | Excluded individuals with high heterozygosity, high relatedness and non-European ancestry. Excluded sample mismatches | | IMPUTE2 | 1000 Genomes version 1 phase 3 |
| B58C | None | >=95% | ≥0.0001 (tested on females only for chromosome X) | ≥1% | Consistent allele frequencies across data deposits (*P*≥0.0001 for pairwise comparisons) and for chrX SNPs, consistent allele frequencies between males and females (*P*≥0.0001). | 500,521 (including 11,696 chrX) | MACH 1.0.18 & Minimac 2012-11-16 | 1000 Genomes Phase 1 March 2012 |
| CROATIA-Korcula | 97% | 98% | 1.00E-06 | 0.01 | | 316,879 | SHAPEIT2, IMPUTE2 | b37; ALL (1000 Genomes Phase 1 integrated release v3, April 2012) |
| CROATIA-Vis | 97% | 98% | 1.00E-06 | 0.01 | | 273,671 | SHAPEIT2, IMPUTE2 | b37; ALL (1000 Genomes Phase 1 integrated release v3, April 2012) |
| ECRHS | | 95% | 0.0001 | 0.01 | High X chromosome heterozygosity for males, sex mismatches, and cryptic relatedness | | MACH 1.0 | HapMap CEU - Build 36, release 22 |
| EPIC population based | None | 95% | 1.00E-08 | Per-plate basis | Monomorphic SNPs; chr 23-26; INDELS; monomorphic; call rate<95%; chr-pos-allels duplicates; delta-AF > 0.2; delta_AF>0.1 if MAF<0.01. Oxford QC:) exclude SNPs if not in HRC ref (no INDEL in HRC ref); 2) exclude if don't match on chr-pos-allele; 3) strand check and flip; 4) exclude if delta-AF>0.2; 5) exclude A/T and G/C SNPs with MAF>0.4 in ref; 6) exclude if chr-pos duplicates | 708,715 | SHAPEIT v2.r790, Oxford | HRC v1.0, 1000 Genomes p3 |

| Study name | Individual call rate filter (applied before imp'n) | SNP call rate filter (applied before imp'n) | SNP HWE *P* filter (applied before imp'n) | SNP MAF filter (applied before imp'n) | Other filters | No of SNPs after filtering (before imp'n) | Imputation software and version | Reference panel used for imp'n |
|---|---|---|---|---|---|---|---|---|
| FinnTwin | Batch1: 98%, Batch2: 95% | Batch1: 97,5%, Batch2: 95% | 1.00E-06 | 0.01 | Genetic ancestry outliers, monomorphic SNPs, high heterozygosity (< -0.03 or >0.05 (method-of-moments F coefficient estimate)), sex mismatches | Batch1: 475637, Batch2: 221814 | Eagle v2.3, Minimac3 v2.0.1, Umich server | HRC r1.1 |
| GS:SFHS | 97% | 98% | 1.00E-06 | 0.01 | Genetic ancestry outliers; monomorphic SNPs; high heterozygosity | 602,451 | SHAPEIT2 v2.r837, Sanger | HRC panel v1.1, European |
| HUNT | 99% | 99% | 0.0001 | 0.01 | Samples that had contamination > 2.5% as estimated with BAF Regress, large chromosomal copy number variants, lower call rate of a technical duplicate pair and twins, gonosomal constellations other than XX and XY, or whose inferred sex contradicted the reported gender, were excluded. Variants were excluded if they were monomorphic, if their probe sequences could not be perfectly mapped to the reference genome, cluster separation was < 0.3, GenTrain score was < 0.15, another assay with higher call rate genotyped the same variant, or they had frequency differences > 15% between data sets | | Michigan server | HRC and custom panel including 2200 HUNT inidividuals with low pass WGS |
| KORA F4 | 0.97 | 0.98 | 5x10-6 | 0.01 | -mismatch of phenotypic and genetic gender<br>- 5s.d. from mean heterozygosity rate<br>- check for European ancestry<br>- check for population outlier | 523,260 (chr 1-26)<br>508,532 (chr 1-22)<br>14,096(chrX-nonPAR)<br>444(chrX-PAR1)<br>58(chrX-PAR2) | SHAPEIT v2, IMPUTE v2.3.0 | 1000g phase1 all (ALL_1000G_phase1integrated_v3_impute_mac1) |
| LBC1936 | 0.95 | 0.98 | ≥0.001 | 0.01 | | 549,692 | minimac 2012-11-16 | 1000 Genomes version 3, cosmopolitan |

| Study name | Individual call rate filter (applied before imp'n) | SNP call rate filter (applied before imp'n) | SNP HWE *P* filter (applied before imp'n) | SNP MAF filter (applied before imp'n) | Other filters | No of SNPs after filtering (before imp'n) | Imputation software and version | Reference panel used for imp'n |
|---|---|---|---|---|---|---|---|---|
| Raine | 0.97 | 0.95 | 0.00000057 | 0.01 | High heterozygosity (h < 0.3), high relatedness (pi=0.1875), sex discrepancies, ATCG ambiguous SNPs | | MACH, minimac | 1000G Phase1 Ver 3 |
| SAPALDIA | 97% | 95% | 1.00E-06 | 0.05 | none | 514,633 (610k quad), 616,660 (OmniExpressExome) | Mach 1.0.16.a, minimac-omp RELEASE STAMP 2012-05-29 (autosomes) & MiniMac RELEASE STAMP 2012-11-16 (chr X) | build37, 1000 Genomes |
| SHIP | 0.92 | 0.95 | 1.00E-04 | None | Genetic ancestry outliers; gender mismatch; pi-hat>0.25; monomorphic SNPS | 760,787 | Eagle v2.3, Michigan | HRC v1.1 reference, European |
| SHIP-TREND | 0.94 | 0.95 | 1.00E-04 | None | Genetic ancestry outliers; gender mismatch; pi-hat>0.25; monomorphic SNPS | 1,691,610 | Eagle v2.3, Michigan | HRC v1.1 reference, European |
| TwinsUK | <0.98 | <97% (SNPs with MAF ≥ 5%) or < 99% (for 1% ≤ MAF < 5%) | 1.00E-06 | 0.01 | Genetic ancestry outliers, heterozygosity across all SNPs ≥2 s.d. from the sample mean, observed pairwise IBD probabilities suggestive of sample identity errors | | Michigan Imputation Server | HRC r1.1 |
| YFS | 0.95 | 0.95 | 1.00E-06 | 0.01 | heterozygosity, relatedness | 546,674 | SHAPEIT v1 and IMPUTE v2.2.2 | 1000 Genomes Phase 1, release v3, March 2012 haplotypes |

**Supplementary Table 4**. Association between rs7697189 and lung function traits in males and females, and genotype-by-sex interaction results on all four lung function traits in males and females combined

| Trait | Lung function males | | Lung function females | | Sex interaction | |
|---|---|---|---|---|---|---|
| | **Beta (SE)** | **P** | **Beta (SE)** | **P** | **Beta (SE)** | **P** |
| $FEV_1$ | 0.052 (0.004) | **2.13E-33** | 0.013 (0.003) | **1.16E-05** | -0.040 (0.005) | **3.15E-15** |
| $FEV_1$/FVC | 0.089 (0.004) | **3.12E-101** | 0.061 (0.003) | **7.49E-76** | -0.028 (0.005) | **8.98E-08** |
| FVC | 0.008 (0.004) | **0.050** | -0.011 (0.003) | **0.0003** | -0.020 (0.005) | **8.71E-05** |
| PEF | 0.072 (0.004) | **7.50E-62** | 0.039 (0.003) | **5.06E-41** | -0.035 (0.005) | **8.78E-12** |

**Supplementary Table 5**. Stratified analyses of the association between rs7697189 and lung function in males and females adjusted for pubertal timing

| Trait | Sex | Without pubertal timing | | Adjusted for pubertal timing | |
|---|---|---|---|---|---|
| | | Beta (SE) | P | Beta (SE) | P |
| $FEV_1$ | Males | 0.0517 (0.0040) | 6.35E-38 | 0.0517 (0.0040) | 4.76E-38 |
| | Females | 0.0219 (0.0035) | 5.97E-10 | 0.0218 (0.0035) | 6.85E-10 |
| $FEV_1$/FVC | Males | 0.0844 (0.0040) | 1.09E-97 | 0.0844 (0.0040) | 1.45E-97 |
| | Females | 0.0645 (0.0035) | 2.93E-74 | 0.0646 (0.0035) | 1.45E-74 |
| FVC | Males | 0.0113 (0.0040) | 0.00521 | 0.0114 (0.0040) | 0.00473 |
| | Females | -0.00708 (0.0036) | 0.0460 | -0.00722 (0.0036) | 0.0419 |
| PEF | Males | 0.0711 (0.0040) | 1.44E-69 | 0.0711 (0.0040) | 1.53E-69 |
| | Females | 0.0586 (0.0035) | 2.70E-61 | 0.0587 (0.0035) | 1.43E-61 |

**Supplementary Table 6**. Association between rs7697189 and *HHIP* expression and rs7697189-by-sex interaction on *HHIP* expression

| *HHIP* probeset ID | Estimate eQTL (SE) | P eQTL | FDR eQTL | Estimate interaction (SE) | P interaction | FDR interaction |
|---|---|---|---|---|---|---|
| X100133899_TGI_at | 0.124 (0.031) | **6.54E-05** | **0.0005** | 0.018 (0.068) | 0.788 | 0.989 |
| X100139086_TGI_at | 0.129 (0.024) | **8.45E-08** | **$2.37 \times 10^{-6}$** | -0.010 (0.050) | 0.838 | 0.989 |
| X100311674_TGI_at | 0.123 (0.027) | **5.61E-06** | **$7.85 \times 10^{-5}$** | -0.009 (0.059) | 0.883 | 0.989 |
| X100148028_TGI_at | 0.143 (0.033) | **1.58E-05** | **0.0001** | -0.004 (0.070) | 0.949 | 0.989 |

**Supplementary Table 7**. Differential expression of *HHIP* in females compared to males

| ***HHIP* probeset ID** | **Estimate (SE)** | **P** |
|---|---|---|
| X100133899_TGI_at | -0.185 (0.044) | **3.14E-05** |
| X100139086_TGI_at | -0.128 (0.033) | **0.000122** |
| X100148028_TGI_at | -0.106 (0.046) | **0.021263** |
| X100311674_TGI_at | -0.173 (0.039) | **6.90E-06** |
